# Tricarboxylic acid cycle and proton gradient in *Pandoravirus massiliensis*: Is it still a virus?

**DOI:** 10.1101/2020.09.21.306415

**Authors:** Sarah Aherfi, Djamal Brahim Belhaouari, Lucile Pinault, Jean-Pierre Baudoin, Philippe Decloquement, Jonatas Abrahao, Philippe Colson, Anthony Levasseur, David C. Lamb, Eric Chabriere, Didier Raoult, Bernard La Scola

**Affiliations:** Aix-Marseille Université, Institut de Recherche pour le Développement (IRD), Assistance Publique - Hôpitaux de Marseille (AP-HM); Microbes, Evolution, Phylogeny and Infection (MEΦI); Institut Hospitalo-Universitaire (IHU) - Méditerranée Infection, 19-21 boulevard Jean Moulin, 13005 Marseille, France; Laboratório de Vírus, Departamento de Microbiologia, Instituto de Ciências Biológicas, Universidade Federal de Minas Gerais, Belo Horizonte, Brazil; Institute of Life Science and School of Medicine, Swansea University, Swansea SA2 8PP, United Kingdom

**Keywords:** giant viruses, Pandoravirus, energy metabolism, ATP production, Lipman system, tricarboxylic acid cycle

## Abstract

Since the discovery of *Acanthamoeba polyphaga* Mimivirus, the first giant virus of amoeba, the historical hallmarks defining a virus have been challenged. Giant virion sizes can reach up to 2.3 µm, making them visible by optical microscopy. They have large genomes of up to 2.5 Mb that encode proteins involved in the translation apparatus. Herein, we investigated possible energy production in *Pandoravirus massiliensis*, the largest of our giant virus collection. MitoTracker and TMRM mitochondrial membrane markers allowed for the detection of a membrane potential in virions that could be abolished by the use of the depolarizing agent CCCP. An attempt to identify enzymes involved in energy metabolism revealed that 8 predicted proteins of *P. massiliensis* exhibited low sequence identities with defined proteins involved in the universal tricarboxylic acid cycle (acetyl Co-A synthase; citrate synthase; aconitase; isocitrate dehydrogenase; α-ketoglutarate decarboxylase; succinate dehydrogenase; fumarase). All 8 viral predicted ORFs were transcribed together during viral replication, mainly at the end of the replication cycle. Two of these proteins were detected in mature viral particles by proteomics. The product of the ORF132, a predicted protein of *P. massiliensis*, cloned and expressed in *Escherichia coli*, provided a functional isocitrate dehydrogenase, a key enzyme of the tricarboxylic acid cycle, which converts isocitrate to α-ketoglutarate. We observed that membrane potential was enhanced by low concentrations of Acetyl-CoA, a regulator of the tricarboxylic acid cycle. Our findings show for the first time that energy production can occur in viruses, namely, pandoraviruses, and the involved enzymes are related to tricarboxylic acid cycle enzymes. The presence of a proton gradient in *P. massiliensis* coupled with the observation of genes of the tricarboxylic acid cycle make this virus a form a life for which it is legitimate to question ‘what is a virus?’.

## Introduction

Since the discovery of *Acanthamoeba polyphaga* Mimivirus (APMV)(1), giant viruses of amoeba have challenged the historical definition and classification of viruses (2). With a virion size larger than 200 nm (1) encompassing genome sizes larger than 250 kb (3), giant viruses differ from all previously described viruses to date. In 2013, *Pandoravirus salinus*, the first Pandoravirus, broke all the viral size records with a genome size of 2.5 Mbp and 1-µm-diameter viral particles (4). Moreover, Pandoravirus genomes do not harbor any gene(s) encoding capsid protein(s), another hallmark of viral biology, and as observed by transmission electron microscopy (TEM), they utilize host cellulose production to build tegument (4, 5). Among later scientific discoveries causing giant viruses to challenge the virus definition were the findings of associated virophages, which depicted for the first time a virus being infected by another virus (6). Ten years later in 2016, the MIMIVIRE system was identified as a mechanism of defense in Mimiviruses against these invading virophages (7). This was the first time that a mechanism for destroying alien DNA, analogous to CRISPR in bacteria, was observed to function in a virus. In 2018, the identification of Mimivirus proteins involved in protein translation again challenged another key feature of the definition of viruses (8). Subsequently, an almost complete protein translation apparatus was discovered in *Tupanvirus* and *Klosneuviruses* (9, 10). More recently, it was found that genes encoding multiple and unique cytochromes P450 monooxygenases commonly occur in giant viruses in the *Mimiviridae, Pandoraviridae*, and other families in the proposed order Megavirales (11, 12).

Moreover, tupanviruses also harbor a gene coding for citrate synthase (13). Recent data indicate that some giant viruses could have genes that are involved in metabolic pathways such as fermentation, sphingolipid biosynthesis and nitrogen metabolism (14). These genes are believed to be used by these viruses to manipulate host metabolic pathways but no evidence suggests that they do not use these gene products themselves for their own metabolism. As giant viruses have challenged most of the criteria for the virus definition, we decided to test another key hallmark of independent life, namely, the ability to produce energy. To test this idea, we used the giant virus *Pandoravirus massiliensis*, which we recently isolated (15). This family of viruses stands uniquely apart from other giant viruses of amoebas because of their huge gene content, with more than 80% of their predicted gene products being ORFans (no homologs in international protein databases). Hence, this virus provides a novel viral system for the discovery of genes with currently unknown functions. In the living world, energy generation is mostly associated with the creation of a proton gradient. Thus, we searched for energy gradients in *P. massiliensis*. We were able to observe the presence of a proton gradient in this virus, and surprisingly, it was mainly present in the mature particles. We then searched for genes that could be associated with this proton gradient. No genes involved in the respiratory chain or with identity to ATP synthase were detected. However, genes having homologies with nearly all enzymes of the tricarboxylic acid (TCA) cycle were observed. These genes were transcribed together, and the product of at least one gene, isocitrate dehydrogenase (IDH), was functional. These findings position this virus as a form a life for which it is legitimate to now ask the question: ‘What is a virus?’

## Materials and methods

### *P. massiliensis* immunofluorescence staining using mouse specific polyclonal antibodies

To avoid confusing virus staining virus with staining of the amoeba mitochondria, we first immunized a mouse with *P. massiliensis* by the subcutaneous route. After three inoculations, mouse serum containing polyclonal antibodies specific to *P. massiliensis* was collected and adsorbed on uninfected *A. castellanii* lysate to remove non-specific antibodies targeting amoeba (16). To permeabilize the cell membranes and saturate the non-specific binding sites, cells were incubated in fetal calf serum with 1% (v/v) Triton X-100 in phosphate-buffered saline (PBS) for 1 h. Infected amoebas were incubated overnight at 28°C in a humidified chamber with anti-Pandoravirus antibodies. Subsequently, the samples were washed three times with 0.1% (v/v) Triton X100 in PBS. Each sample was incubated with fluorescein isothiocyanate (FITC)-conjugated goat anti-mouse IgG (Immunotech, Marseille, France) for 60 min at 28°C in a humidified chamber and finally washed three times with PBS.

### Assessment of a membrane electrogradient in *P. massiliensis* virions

Amoeba were infected with *P. massiliensis* at a multiplicity of infection (MOI) of 10 in IBIDI^®^ petri µ-dishes. Each sample was dyed with MitoTracker Deep Red 633 (Invitrogen, Carlsbad, California, USA), blocked with acetone, and washed three times with PBS. To target a proton gradient potential difference across the *P. massiliensis* particle membranes, two reagents were used: MitoTracker Deep Red 633 and tetramethyl rhodamine (TMRM) reagent (Thermo Fisher Scientific). For each reagent, *P. massiliensis* viral particles freshly released from 24-h infected ameba cultures were used. Briefly, lysed amoeba infected by *P. massiliensis* were centrifuged 10 min at 500 × *g*, and cellular debris was discarded. The viral supernatant was centrifuged 20 min at 6800 × *g*, and the pellet was resuspended twice in PAS. The second time, the viral pellet was resuspended in survival buffer. In control experiments, sample cultures of *Staphylococcus aureus* were used as positive controls, and viral supernatant from cowpoxvirus cultured on Vero (ATCC CCL-81) African green monkey kidney cells (17) was used as negative control. The MitoTracker Deep Red 633 (50 µg) was reconstituted in 1.5 mL of Peptone Yeast culture medium (PYG) culture medium to obtain a solution stock of 33.3 µg/mL, which was subsequently tested in survival buffer in IBIDI^®^ petri µ-dishes previously coated with poly-L lysine to retain adherent cells, even at late time points of infection and after the wash steps. MitoTracker was added to the samples 1 h post-infection, and images recorded 2, 4, and 6 h post-infection. For late time points of the viral cycle (8 h, 10 h, 12 h, 14 h, 16 h), MitoTracker was added to the samples 7 h post-infection. The medium was replaced with PYG. Subsequently, 34 µL of MitoTracker Deep Red 633 pre-incubated at 37°C was added to each petri µ-dish and incubated for 45 min at 37°C. Each sample was washed to remove the excess fluorescent dye, and 2 ml of survival buffer was added. For TMRM, a 1-ml volume of viral particles was deposited in a petri µ-dish, after which 1 µl of stock solution of TMRM (100 µM) was directly added and incubated 30 min at 30°C.

### Assessment of the effect of the decoupling agent CCCP on viral particles

The effect of carbonyl cyanide m-chlorophenylhydrazone (CCCP) (Sigma Aldrich C2759; Saint-Louis, Missouri, USA), an inhibitor of oxidative phosphorylation that acts by dissipating the electrochemical gradient induced by the proton concentration, was assessed on *P. massiliensis* virions treated with TMRM. CCCP reagent (100 μM, 200 μM, 300 μM, 400 µM) was directly added in tubes containing ≈ 10^7^ viral particles /ml. Negative control were *P. massiliensis* virions without CCCP. Samples were incubated at 35°C overnight and then transferred to petri µ-dishes. Next, 1 µl of a stock solution of TMRM (100 µM) was added and incubated 30 min at 30°C. Images were acquired by confocal microscopy using a Zeiss LSM 800 microscope. The infectivity of *P. massiliensis* particles was assessed before and after incubation with CCCP by calculating the TCID50 using the method of Reed and Muench (18). The potential impact of CCCP on viral replication cycle was also assessed by immunofluorescence and qPCR. *A. castellanii* strain Neff cells were inoculated with viral particles previously incubated with 400 µM CCCP and washed three times. As at the end of the cycle, no difference was observed, so we focused on early time points of the viral cycle. Infected amoeba were collected 45 min (H0) and 3h (H3) post-infection. *P. massiliensis* particles spotted on slides were labeled with anti-*P. massiliensis* specific antibodies according to the protocol described above. qPCR was carried out using DNA from the collected cells with a system targeting the DNA polymerase gene of *P. massiliensis* (forward primer: 5′-ATGGCGCCCGTCTGGAAG; reverse primer: 5′-GGCGCCAAAGTGGTGCGA). qPCR was performed with a LightCycler® 480 SYBR Green 1 Master reaction mix (Roche Diagnostics, Mannheim, Germany), following the manufacturer’s temperature program with 60°C for the primer hybridization and elongation temperature.

### Bioinformatics analyses

The *P. massiliensis* genome was analyzed by BLASTp analyses against the GenBank nr database using an e-value threshold of 1 ×10^−2^. The search for enzymes potentially involved in energy metabolism was performed by delta BLAST analyses against the Conserved Domain Database (19, 20). For some predicted ORFs having hits with low similarity, PSI Blast, HHPRED analyses (21) and structure prediction using the PHYRE2 server were performed (22). Orthologs in other pandoravirus genomes were searched using the ProteinOrtho tool with a 30% identity percentage threshold and 50% as a coverage percentage threshold **(23)**. Gene products of all pandoraviruses were also analyzed by BLASTp against the COG database (24, 25). The viral ORFs harboring a hit against class C of the COG (energy metabolism) with a bitscore >50 were considered statistically significant.

### Transcriptome sequencing (RNA-seq) on *P. massiliensis*

*P. massiliensis* replicates in *Acanthamoeba castellanii* Neff (ATCC 30010). The transcriptome of *P. massiliensis* strain BZ81 was assessed as previously described using amoebas infected by *P. massiliensis* as well as freshly released mature viral particles (15). Mature virions were collected 11 h following amoeba inoculation, passed through 5-µm-pore filters, and centrifuged at 500 × *g* for 10 min to remove all amoeba debris.

### qRT-PCR of suspected *P. massiliensis* TCA cycle genes

Viral DNA isolated from 200 µL of viral supernatant from *P. massiliensis* culture was extracted using the EZ1 tissue kit (Qiagen, France) according to the manufacturer’s recommendations. RNA was extracted using the RNeasy mini kit (Qiagen, France) at different time points of the *P. massiliensis* replication cycle, from H0 (i.e., 45 min after infection of ameba cells by viral particles) until H16 post-infection (release of neo-synthetized virions), according to the manufacturer’s recommendations.).

Total RNA was reverse-transcribed into cDNA using the SuperScript VILO Synthesis Kit (Invitrogen, France). Nucleotide primers targeting the 7 selected ORFs of the *P. massiliensis* gene sequences were designed using the primer3 tool (26) (Table 1). qPCR was carried out using LightCycler® 480 SYBR Green 1 Master reaction mix (Roche Diagnostics, Mannheim, Germany) following the manufacturer’s temperature program with 62°C as the primer hybridization and elongation temperature. Each experiment was performed in triplicate. The results were considered positive if the cycle threshold obtained in three replicates was less than 35.

**Table 1.**
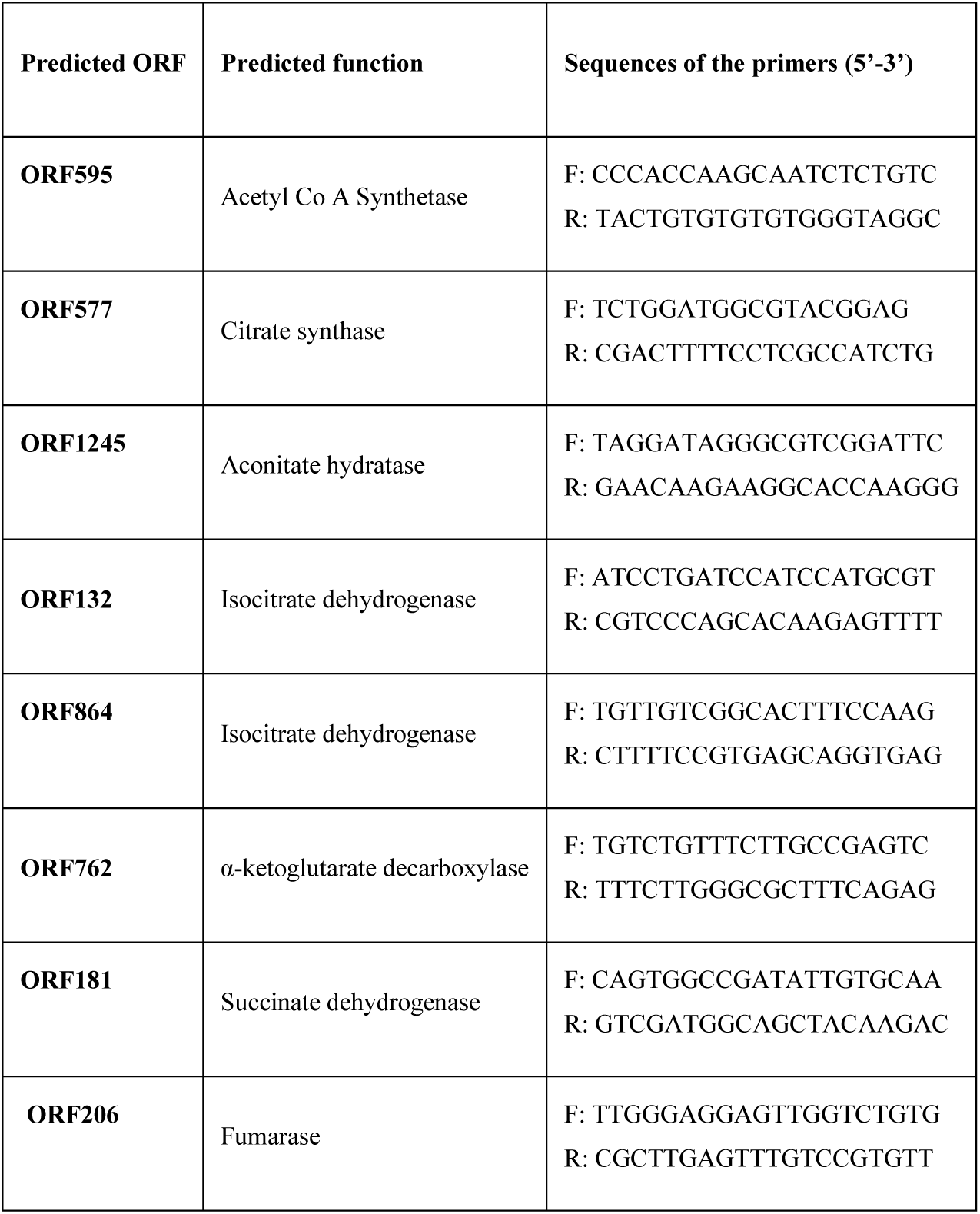
qRT-PCR primers used in the present study of the predicted ORFs of *P. massiliensis*.

### Proteome analysis of *P. massiliensis*

Protein extraction was carried out on purified viral particles and amoebas infected with *P. massiliensis* at different time points of the replication cycle (H0 to H16). Briefly, samples were rapidly lysed in dithiothreitol (DTT) solubilization buffer (2% (w/v) SDS, 40 mM Tris– HCl, pH 8.0, 60 mM DTT) with brief sonication. The 2D Clean-Up kit was used to eliminate nucleic acids, salts, lipids, and other molecules that were not compatible with immunoelectrophoresis. Next, 1D gel electrophoresis analysis was performed with Ettan IPGphor II control software (GE Healthcare). For 2D gel electrophoresis, buffer (50 mM Tris-HCl, pH 8.8, 6 M urea), 30% (v/v) glycerol, 65 mM DTT reducing solution, alkylating solution of iodoacetamide at 100 mM, and an SDS-PAGE gel with 12% (v/v) acrylamide were used. Protein migration was performed under a constant electric field of 25 mA for 15 min, followed by 30 mA for ≈5 h. Silver nitrate was used for protein staining. Proteins of interest were excised from the gel and analyzed.

For global proteomic analysis, the protein-containing solution was subjected to dialysis and trypsin digestion. Dialysis was carried out twice using Slide-ALyzer 2K MWCO dialysis cassettes (Pierce Biotechnology, Rockford, IL, United States) against a solution of 1 M urea and 50 mM ammonium bicarbonate pH 7.4: 4 h and overnight. Protein digestion was carried out by adding 2 μg of trypsin solution (Promega, Charbonnières, France) to the alkylated proteins, followed by incubation at 37°C overnight. Digested protein samples were desalted using detergent columns (Thermo Fisher Scientific, Illkirch, France) and analyzed by mass spectrometry on a Synapt G2Si Q-TOF traveling wave mobility spectrometer (Waters, Guyancourt, France) as described previously (27). An internal protein sequence database built primarily with two types of amino acid sequences was used as follows: (i) sequences obtained by translating *P. massiliensis* ORF (ii) sequences obtained by translating the whole genome into the six reading frames and then fragmenting the six translation products into 250-amino-acid-long sequences with a sliding step of 30 amino acids. Contiguous sequences that were positive for peptide detection were fused and reanalyzed.

### Cloning, expression and purification of predicted *P. massiliensis* TCA cycle enzymes

*P. massiliensis* genes encoding the predicted TCA enzymes (ORFs 132, 181, 206, 577, 595, 762, 864, 1245) were designed to include a Strep-tag at the N-terminus and optimized for *Escherichia coli* expression. Genes were synthesized by GenScript (Piscataway, NJ, USA) and ligated between the NdeI and NotI sites of a pET22b(+) plasmid. Competent BL21(DE3) cells grown in autoinducing ZYP-5052 medium were used for expression of the recombinant proteins. To produce each protein, the culture was shaken at 37°C until an O.D.600 nm of 0.6 was reached, after which the temperature was lowered to 20°C for 20 h. Cells were harvested by centrifugation (5 000 × g, 30 min, 4°C), and the resulting pellet was resuspended in 50 mM Tris pH 8, 300 mM NaCl and stored at −80°C overnight. The crude extract was thawed and incubated on ice for 1 h following the addition of lysozyme, DNAse I and phenylmethylsulfonyl fluoride (PMSF) to final concentrations respectively of 0.25 mg/mL, 10 µg/mL and 0.1 mM. Partially lysed cells were sonicated using a Q700 sonicator system (QSonica), and cell debris was removed following a centrifugation step (12 000 g, 20 min, 4°C). Proteins of interest were purified with an ÄKTA avant system (GE Healthcare) using strep-tag affinity chromatography (Wash buffer: 50 mM Tris pH 8, 300 mM NaCl and Elution buffer: 50 mM Tris pH 8, 300 mM NaCl, 2.5 mM desthiobiotin) on a 5-mL StrepTrap HP column (GE Healthcare). Recombinant protein expression was confirmed by MALDI-TOF MS analysis of excised gel bands previously isolated by SDS-PAGE. Protein concentrations were measured using a Nanodrop 2000c spectrophotometer (Thermo Scientific).

### IDH activity assay and kinetics

IDH activity assays were performed using the Isocitrate Dehydrogenase Activity Assay kit (MAK062) from Sigma-Aldrich (St. Louis, MS, USA) and monitored with a Synergy HT microplate reader (BioTek, Winooski, VT, USA). Reactions were carried out in duplicate at 37°C in a 96-well plate containing a final volume of 100 µL in each well. Conversion of the isocitrate substrate to α-ketoglutarate was monitored for 30 min following absorbance variations at 450 nm, corresponding to the production of NADH. A NADH standard curve was plotted and allowed quantification of the produced NADH with our enzyme and calculation of its specific activity. Initial velocities were calculated using Gen5.1 software (BioTek), and the obtained mean values were fitted using the Michaelis-Menten equation in Prism 6 (GraphPad Software, San Diego, CA, USA).

### Assessment of the effect of acetyl CoA on *P. massiliensis* viral particles

Acetyl-CoA (Sigma-Aldrich) was added at concentrations of 0.8 mM, 0.4 mM, 0.2 mM, 0.1 mM, 0.01 mM, and 0.001 mM to 1 mL of the viral suspension of purified particles (≈ 10^8^ particles/ml). The negative control consisted of viral particles without acetyl CoA. The samples were incubated at 30°C for 24 h. After incubation, the samples were transferred into petri µ-dishes and stained with TMRM by following the above-described protocol. Images were acquired using an LSM 800 confocal microscope. Image processing and fluorescence intensity evaluations were conducted using Zen Bleu software.

### Statistical analysis

Statistical analysis was performed using GraphPad Prism for Windows. Statistical differences were evaluated by one-way ANOVA. Statistical significance was set at p <0.05.

## Results

### Detection of membrane potential in *P. massiliensis* virions

Membrane potential in *P. massiliensis* virions was assessed during the replication cycle in *A. castellanii* and mature virions freshly released from amoebas. During the viral cycle of *P. massiliensis* in *A. castellanii*, a variable proportion of MitoTracker-labeled viruses was observed (Figure1). The specificity of the labeling was ensured by the co-localization of FITC-conjugated anti-*P. massiliensis* antibodies.

**Figure 1.**
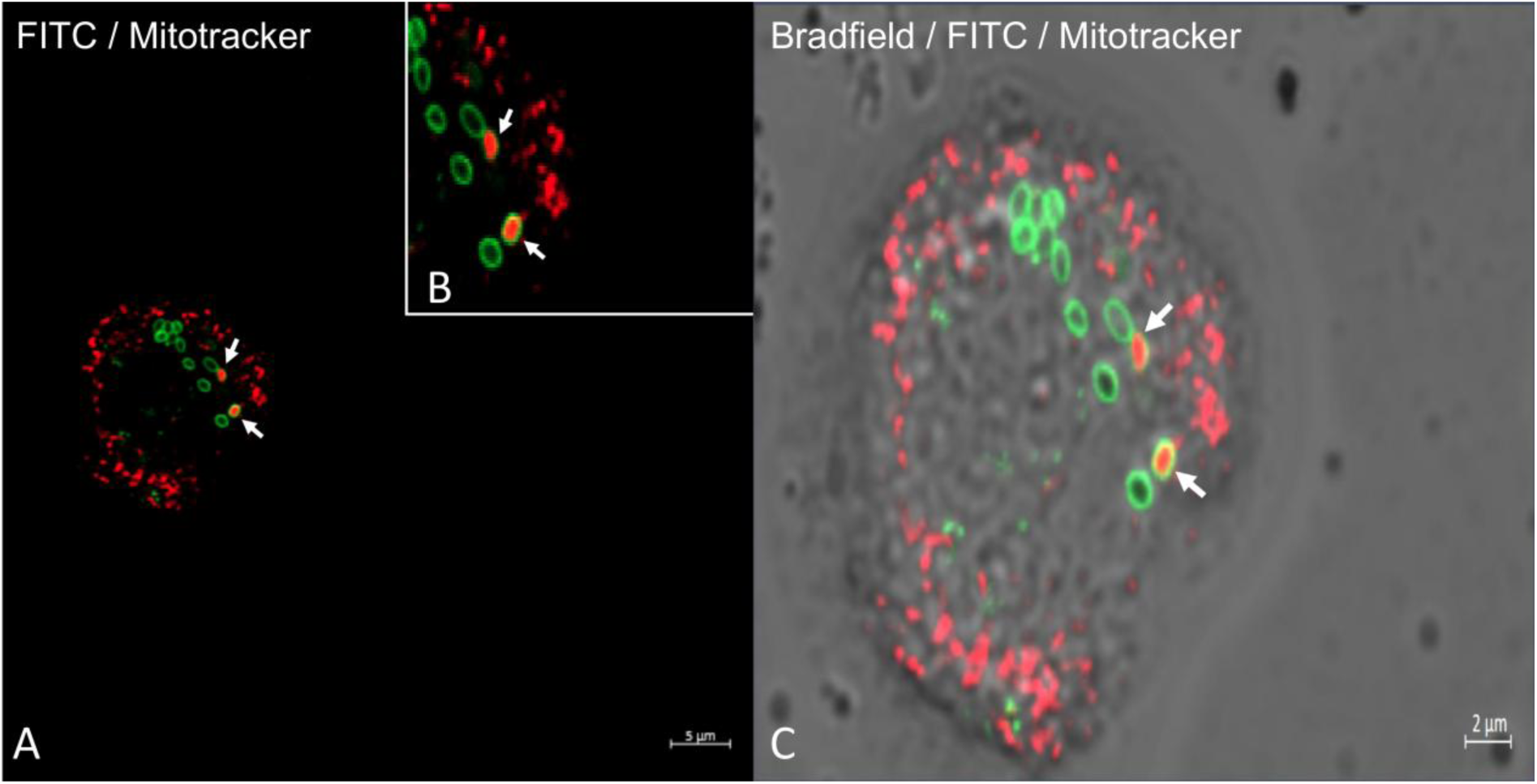
Confocal imaging of amoeba infected by *P. massiliensis* stained with MitoTracker Deep Red (in red) and with specific anti-*P. massiliensis* antibodies (in green). A,B Colocalization of the MitoTracker signal (in red) with virus marked by specific antibodies (FITC) (arrows). C: Merge of Bradfield, FTIC and MitoTracker fluorescence.

An analogous experiment performed using mature viral particles showed that approximately 20% of the total number of particles was labeled both by anti-*P. massiliensis* specific antibodies and MitoTracker Deep Red633 (Figure 2). Moreover, the viral particles were also marked by TMRM staining (under the TRITC wavelength (532 nm), with a fluorescent signal (Figures 3,4)), similar to the results obtained for the *S. aureus* positive control. No fluorescence was observed in the cowpoxvirus negative control experiments.

**Figure 2.**
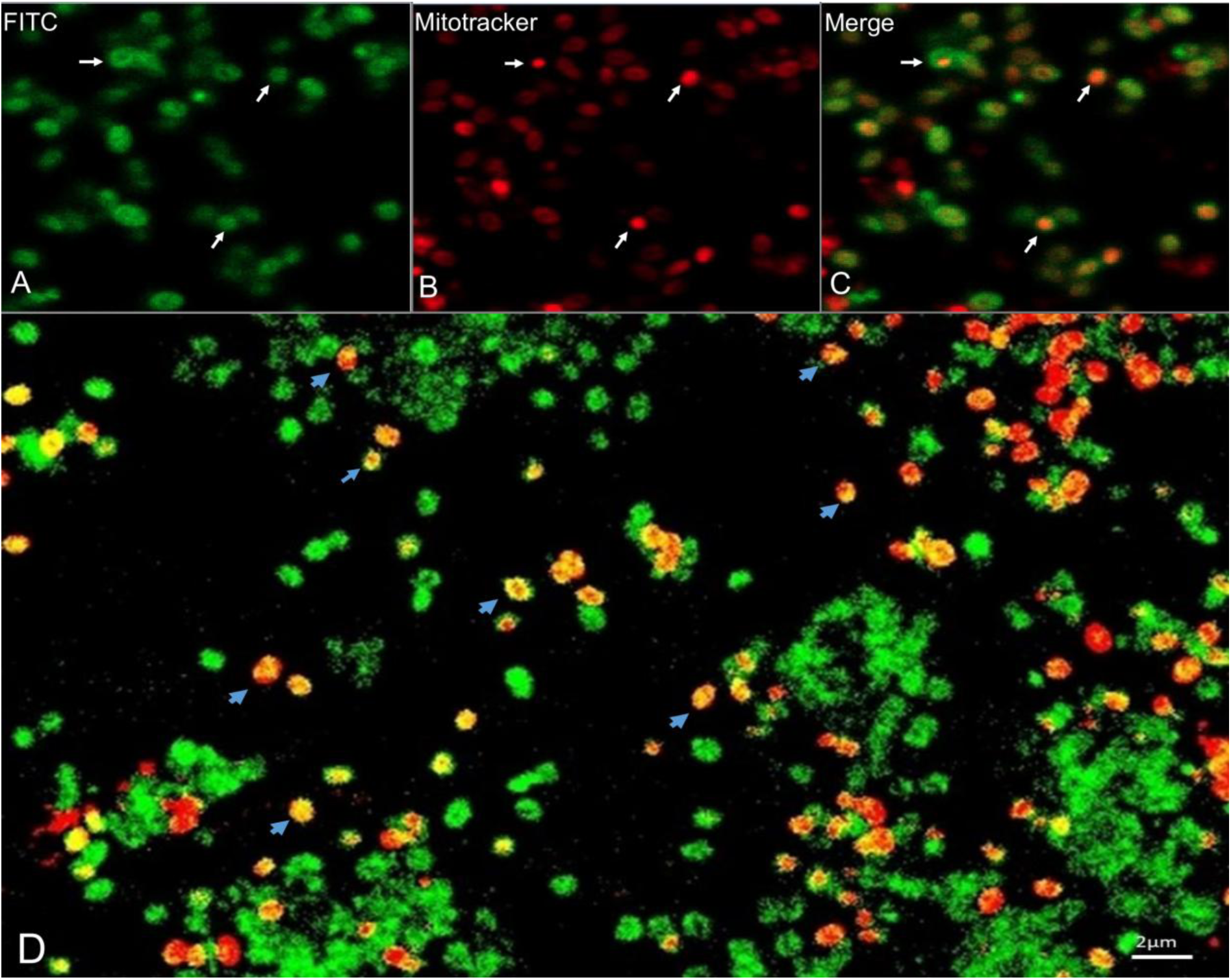
Confocal imaging of MitoTracker staining of viral mature particles of *P. massiliensis*. A: Specific antibody stained viral particles (FITC) (white arrows). B: MitoTracker Deep Red (red) incorporated into *P. massiliensis* particles (white arrow). C,D: Colocalization of the MitoTracker signal (red) with *P. massiliensis* virions marked by specific antibodies (white and blue arrow).

**Figure 3.**
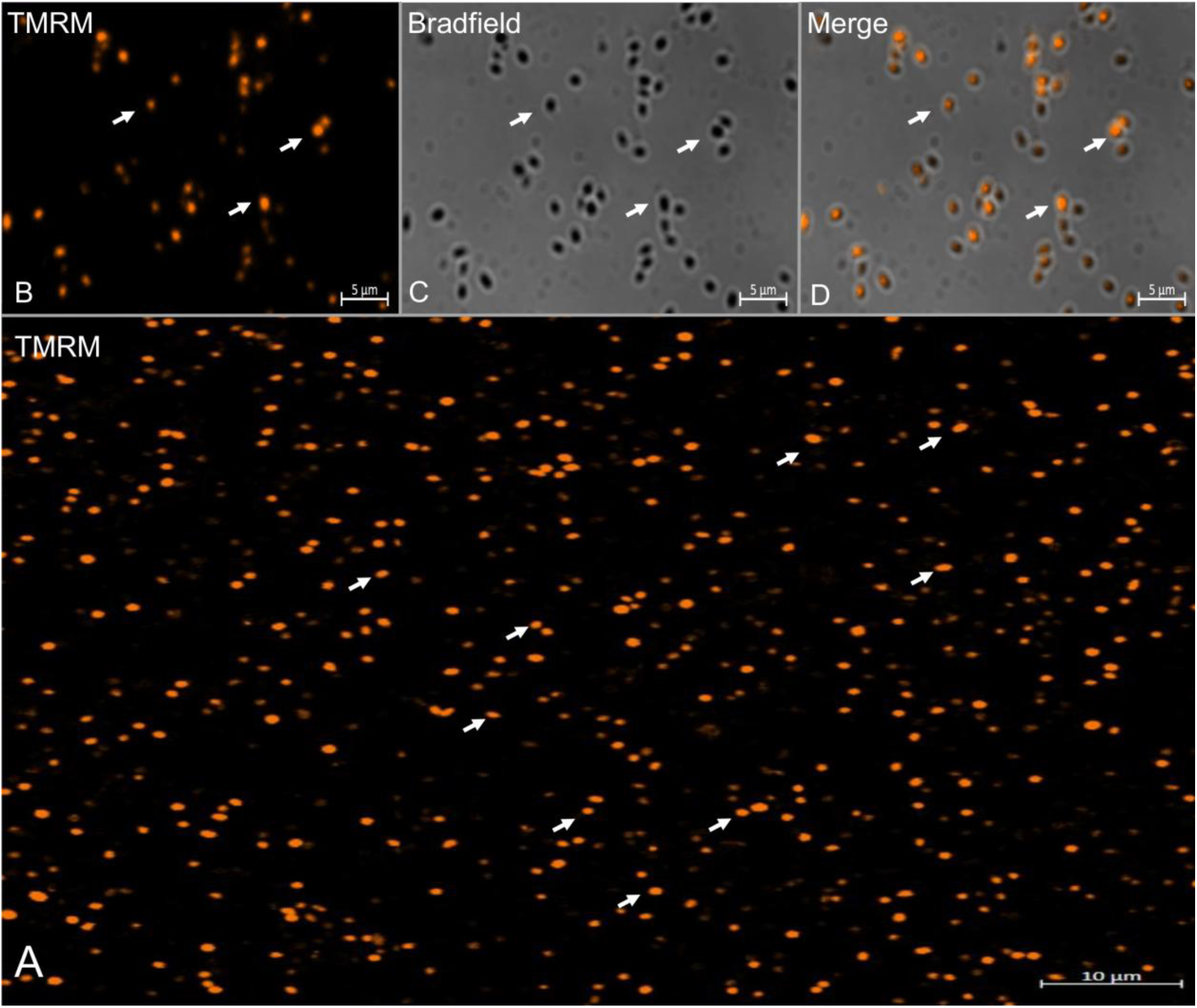
Confocal imaging of TMRM staining of purified *P. massiliensis* virions. A,B: Viral mature particles stained with TMRM (arrows). C: Bradfield channel. D: Merge of Bradfield and TMRM showing the internalization of the TMRM signal in viral particles.

**Figure 4.**
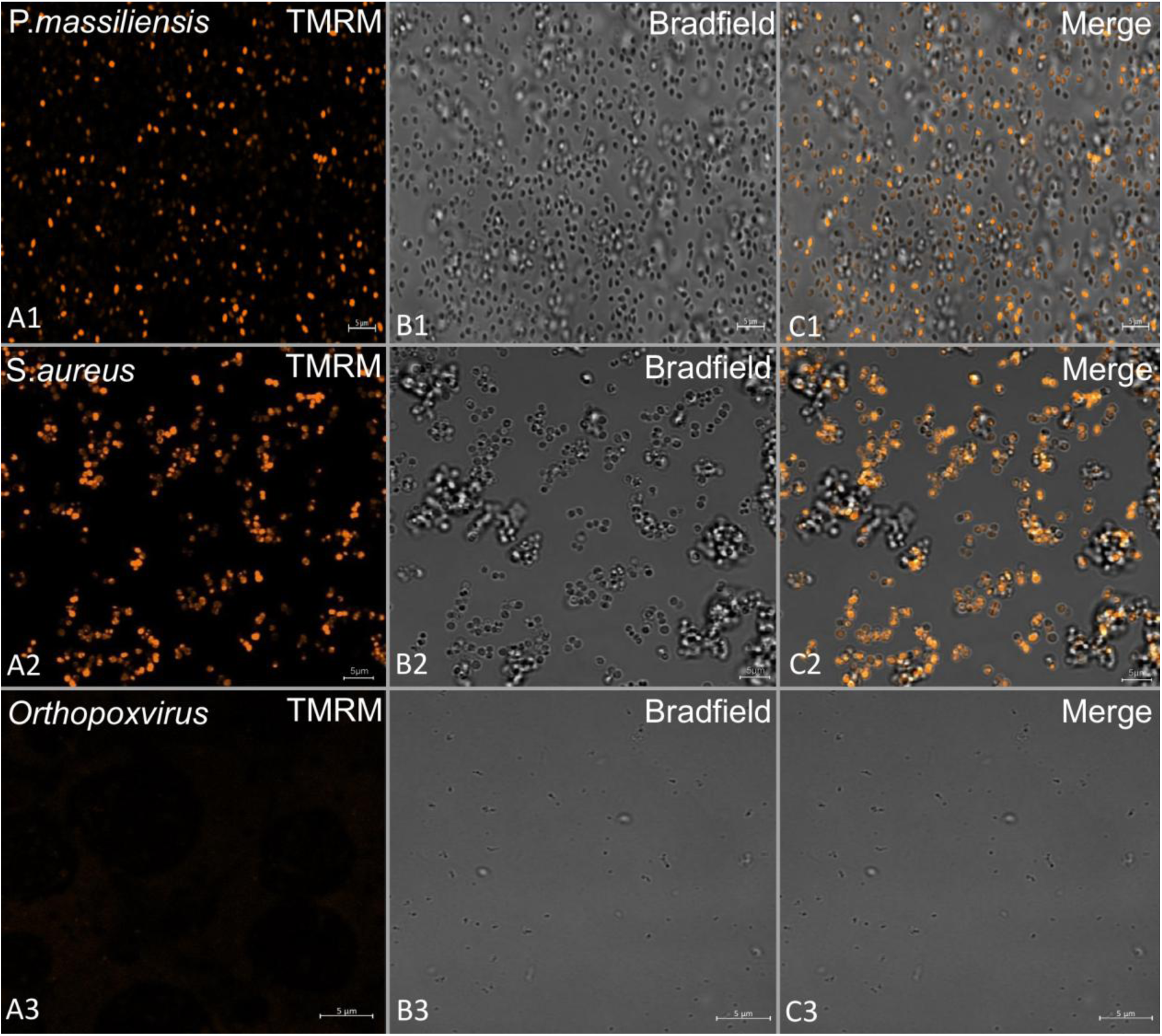
TMRM fluorescent staining of viral mature particles of *P. massiliensis*. (A1-C1). Positive control consisting of *S. aureus* (A2-C2) and negative controls consisting of cowpoxvirus (A3-C3) are shown.

### Viral particles treated with CCCP

*P. massiliensis* particles were incubated with a range of concentrations of CCCP: 100 µM, 200 µM, 300 µM, and 400 µM. For the *S. aureus* positive control, the fluorescent signal generated by TMRM decreased significantly (p<0.05) in the presence of all concentrations of CCCP compared with the negative controls (viral particles without CCCP) (Figure 5).

**Figure 5.**
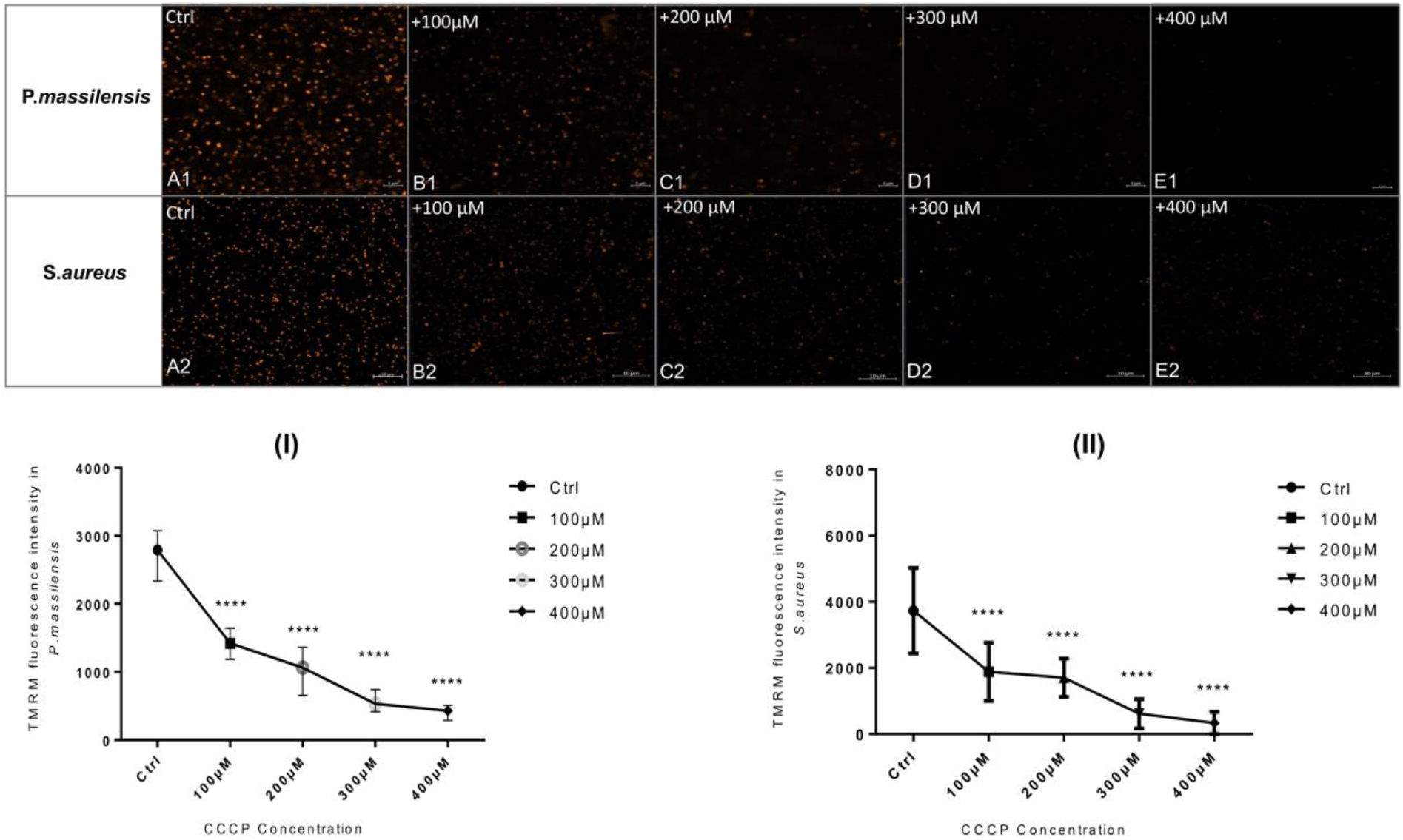
Evaluation of the fluorescence intensity of TMRM after CCCP treatment. (A1-E1): Confocal imaging of TMRM staining following CCCP treatment of *P. massiliensis* particles. A1: Control experiment using untreated *P. massiliensis* particles. B1,E1: Confocal imaging of *P. massiliensis* virions treated with different concentrations of CCCP. (A2-E2): Confocal imaging of TMRM staining after CCCP treatment of the positive control (*S. aureus*). A2: Control experiment showing untreated *S. aureus*. B2,E2: *S. aureus* treatment with a different concentration of CCCP. (I): Estimation of TMRM fluorescence intensity of *P. massiliensis* particles following CCCP treatment. (II): Estimation of TMRM fluorescence intensity of *S. aureus* following CCCP treatment.

The titer of these viral particles after incubation with CCCP did not show any significant difference compared with the negative control (10^7,2^ TCID_50_/ml). Delta-Ct between the negative control (CCCP untreated samples) and viral particles preincubated with CCCP at 400 µM was 1.85 and 2.57 for the H0 and H3 post-infection time points, respectively. Immunofluorescence revealed a smaller number of labeled viral particles preincubated with CCCP at 400 µM on amoeba cells (1313 and 1613 at H0 and H3, respectively) than the negative control (1658 and 1889 at H0 and H3, respectively), but the difference was not significant (Figure 6).

**Figure 6:**
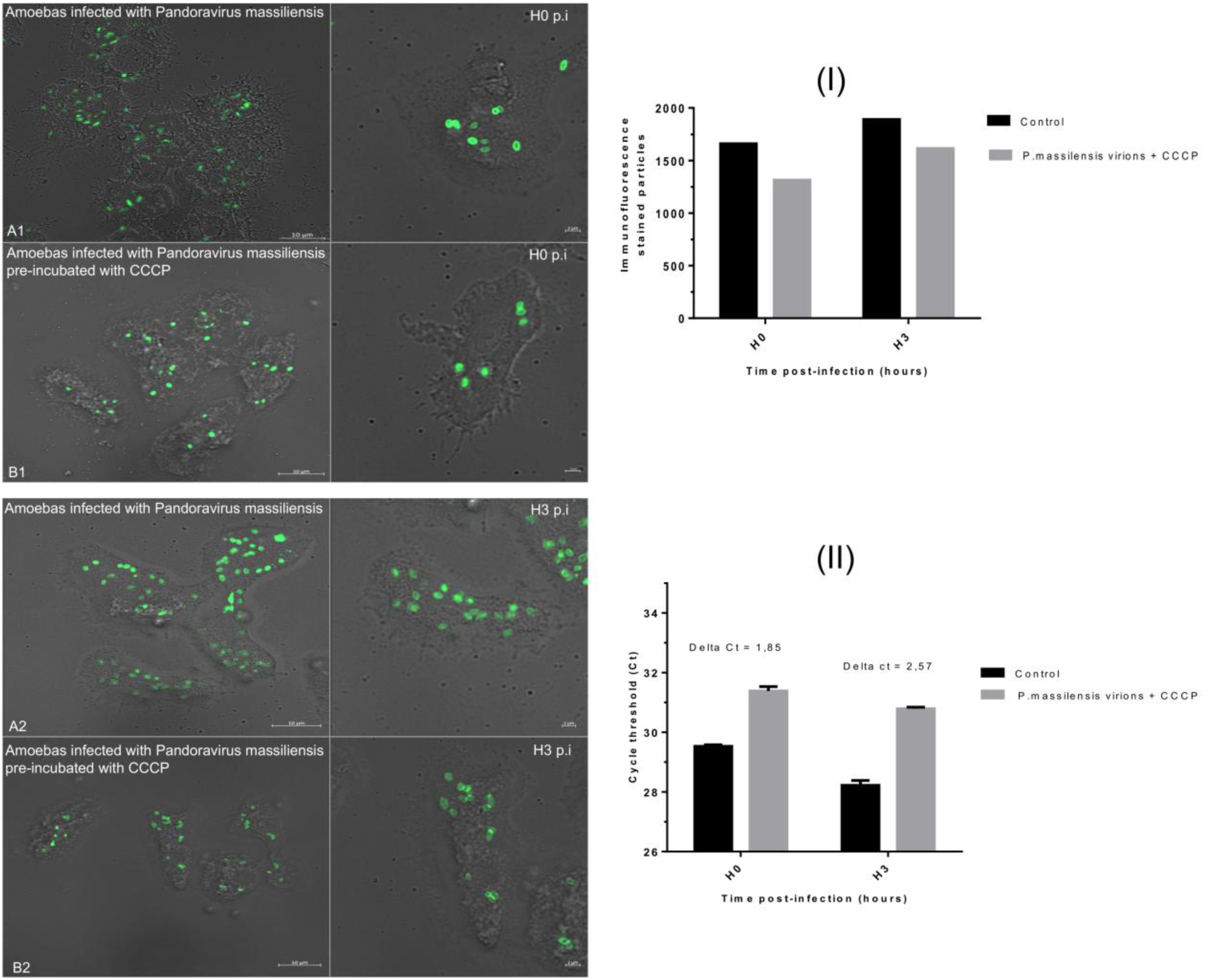
Assessment of the CCCP treatment effect on *P. massiliensis* infectivity. (A1-B2): Immunofluorescence confocal imaging of stained amoeba infected with *P. massiliensis* particles preincubated with and without CCCP. (A1) Negative control: *P. massiliensis* particles (green) in amoeba at H0 p.i. (B1) *P. massiliensis* + CCCP in amoeba at H0 p.i. (A2) Negative control: *P. massiliensis* virions in the absence of CCCP (green) in amoeba at H3 p.i. (B2) *P. massiliensis* particles (green) in amoeba at H3 p.i. (I): Estimation of the number of stained particles of *P. massiliensis* particles without and with the highest concentration of CCCP (400 µm) per/100 amoebas at H0, H3 p.i. (II): Representation of the mean threshold cycle (Ct) of the qPCR experiments (triplicate) for isolated *P. massiliensis* DNA before and after CCCP treatment according to the postinfection time from 0 to 3 h.

### Evaluation of the fluorescence intensity of TMRM after incubation of *P. massiliensis* particles with acetyl Co-A

In comparison to the negative control (untreated pandoravirus particles) (Figure 8.A1), the TMRM fluorescent signal significantly increased in the presence of low concentrations of acetyl-CoA (0.01 mM) (figure 8.F1.I) (p<0.05) and significantly decreased in the presence of high concentrations of acetyl CoA (0.8 mM, 0.4 mM) (Figure 7.B1.C1.I) (p <0.05).

**Figure 7:**
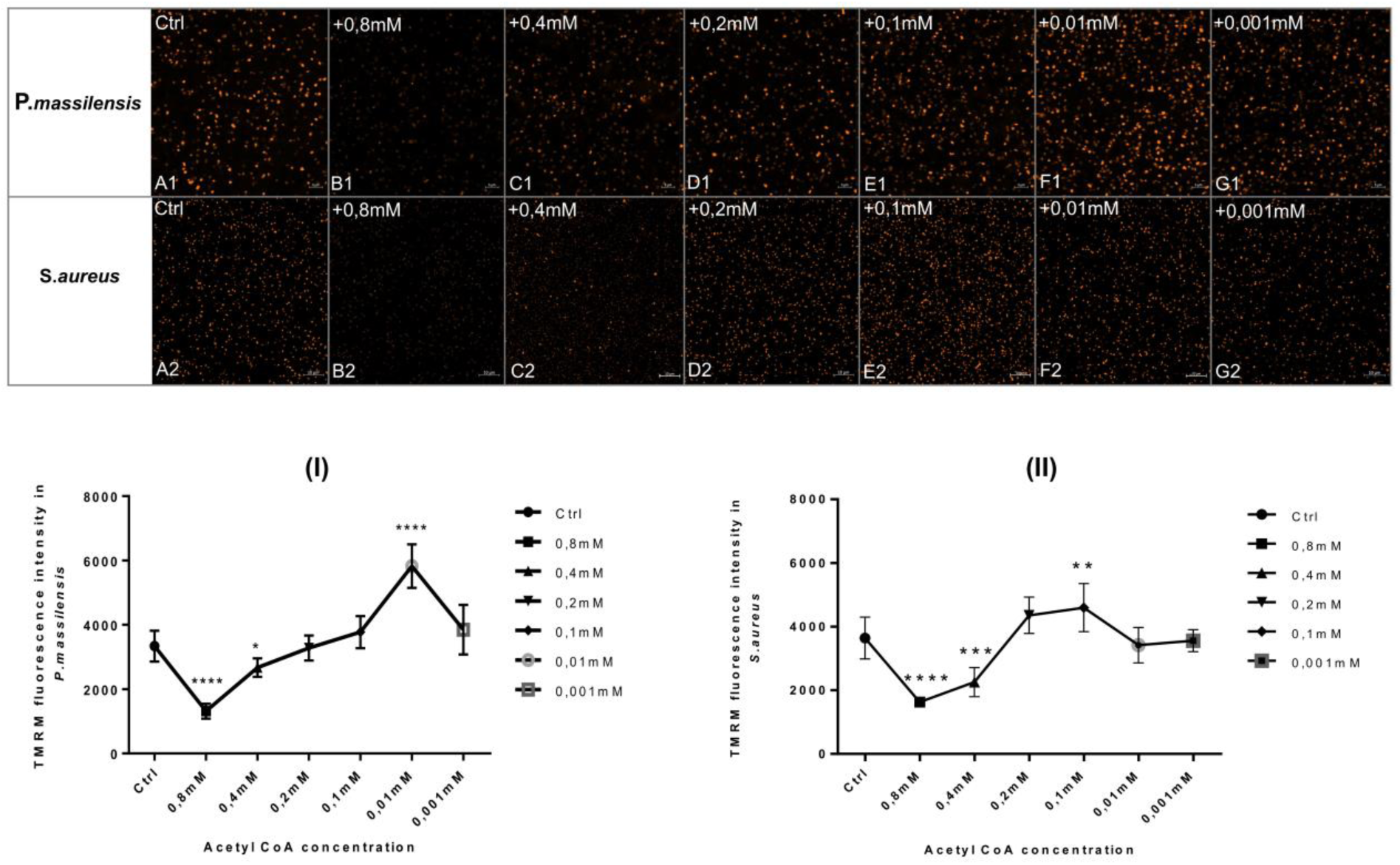
TMRM fluorescence intensity evaluation following acetyl CoA treatment. (A1-G1): Confocal imaging of TMRM staining following acetyl CoA treatment of *P. massiliensis* particles. A1: Control condition with untreated *P. massiliensis* particles. B1,G1: *P. massiliensis* virions treated with different concentrations of acetyl CoA. (A2-G2): Confocal imaging of TMRM staining after acetyl CoA treatment of the positive control (*S. aureus*). A2: Control experiment with untreated *S. aureus*. B2,G2: *S. aureus* treatment with a different concentration of acetyl CoA. (I): Estimation of the TMRM fluorescence intensity of *P. massiliensis* particles after acetyl CoA treatment. (II): Estimation of the TMRM fluorescence intensity of *S. aureus* after acetyl CoA treatment.

**Figure 8:**
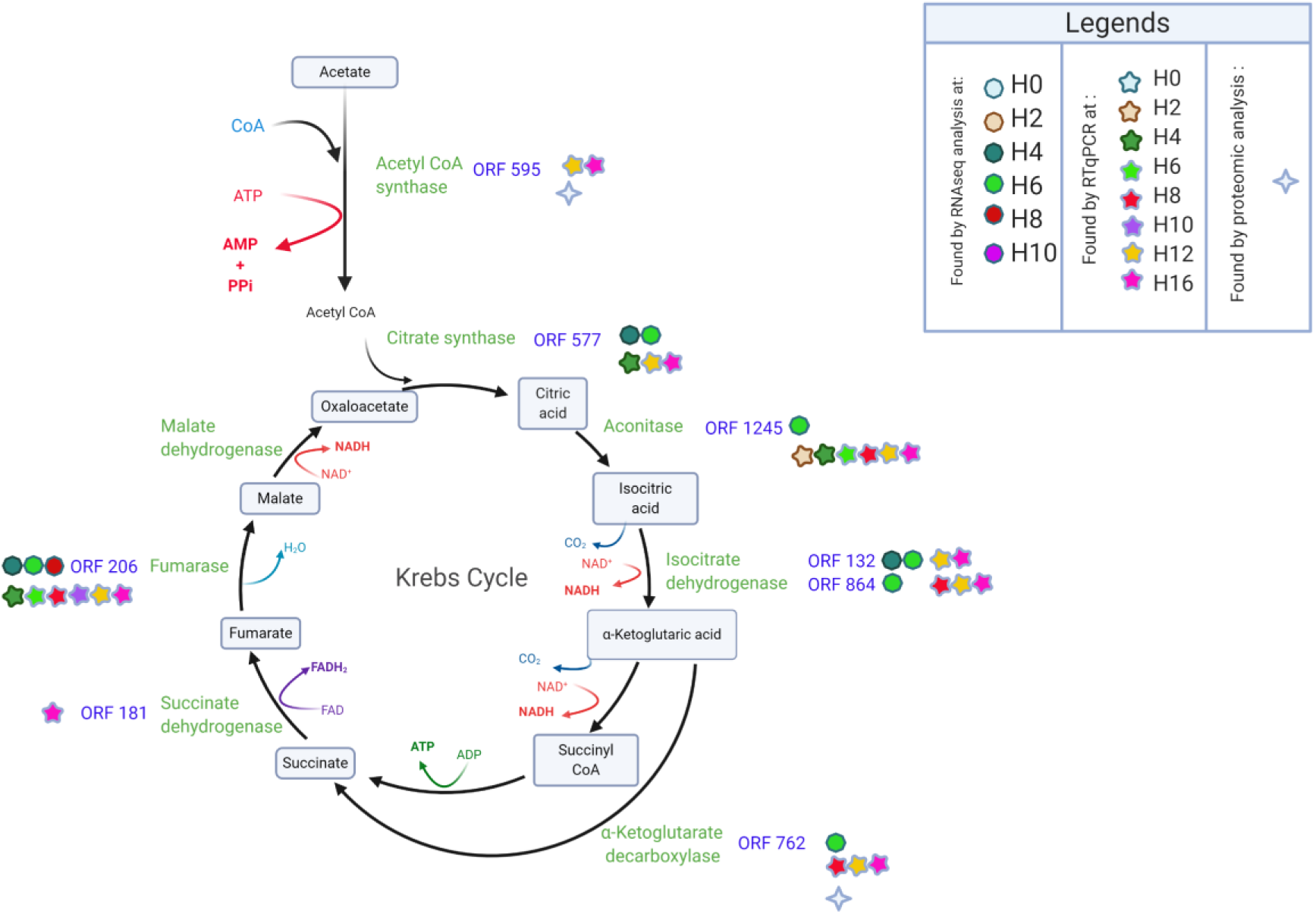
Schematic representation of the TCA cycle showing the predicted ORFs of *P. massiliensis* with similarities to TCA cycle enzymes, and a summary of the results provided by qRT-PCR, RNA sequencing and proteomics.

In positive control experiments using *S. aureus*, the TMRM fluorescent signal increased significantly at low concentrations of acetyl-CoA (0.1 mM) (Figure 7.E2.II) (p<0.05) and decreased significantly in the presence of high concentrations of acetyl CoA (0.8 mM, 0.4 mM) (Figure7.B2.C2.II, Figure8.B2.C2.II) (p <0.05).

### Bioinformatics analyses

Using DELTA-BLAST analyses against the Conserved Domain Database (CDD) (20), low sequence similarity with enzymes involved in the TCA were found. Before concluding that this similarity was not significant, we searched for other predicted *P. massiliensis* gene products with similarities to other enzymes of the TCA cycle (i.e., citrate synthase, aconitase, α-ketoglutarate dehydrogenase, succinyl CoA synthetase, succinate dehydrogenase, fumarase). Low similarities were found for 6 *P. massiliensis* predicted gene products with 6 enzymes of the TCA cycle. The product of ORF577 exhibited 33% identity to the conserved domain PRK05614 of citrate synthase (bitscore 58). A similarity of the ORF1245 gene product was found for domain pfam05681 of aconitase. The *P. massiliensis* ORF132 gene product harbored similarity to isocitrate/isopropyl malate dehydrogenase (COG0473) with a bitscore of 67 and 30% identity. Using a HHPRED with the TIGR PFAM database also revealed a low similarity of ORF132 with TIGR00169, an NAD or NADP dehydrogenase, including dimeric forms of IDH. In addition to the similarity found for ORF132, ORF864 harbored 50% identity to another domain of IDH (pfam03971) (bitscore: 54). No similarity was found for α-ketoglutarate deshydrogenase or succinate thiokinase. However, a low similarity was found for the *P. massiliensis* predicted ORF762 gene product (bitscore: 55; identity: 41%) with alpha-ketoglutarate decarboxylase, which converts alpha-ketoglutarate to succinate, switching the 2 steps of alpha-ketoglutarate dehydrogenase and succinate thiokinase. The ORF181 gene product was approximately 30% identical to domain PRK09078 of the succinate dehydrogenase (bitscore: 77). Finally, domain PRK06246 of fumarase showed 30% identity to the predicted ORF206 gene product (bitscore: 58). The search for a similarity of structure of these 7 ORFs with Phyre2 was inconclusive. No similarity was found for malate dehydrogenase. Of note, a hit with acetyl-CoA synthetase, the immediate step upstream of the first step of the TCA cycle (synthesis of citrate starting from acetyl-CoA) was found for the ORF595 gene product (bitscore: 58; identity: 24%).

BLASTp analyses of these 8 *P. massiliensis* ORFs putatively involved in the TCA cycle against the nr database revealed the predicted enzymatic function for only the gene product of ORF595, with only one hit annotated as acetyl CoA synthetase of *Phalacrocorax carbo*, with 33% identity. An ortholog in other pandoraviruses was also found for the ORF132, ORF181 and ORF206 of *P. massiliensis*. ORF132 was orthologous to YP00948512.1 from *P. neocaledonia*. Orthologs for ORF181 were found in all the other pandoraviruses: cds786 from *P. macleodensis*; cds867 from *P. neocaledonia*; cds120 from *P. salinus*; cds1076 from *P. celtis*; cds1057 from *P. quercus*; pi_168 from *P. inopinatum* (YP009119137.1); and cds943 from *P. dulcis*, with e-values and identities ranging from 1.52e-52 to 5.29e-56 and 74 to 46%, respectively. Orthologs for ORF206 were found in *P. neocaledonia* (cds851), *P. macleodensis* (cds769), *P. dulcis* (cds1004), *P. salinus* (cds1274), and *P. quercus* (cds1120). Stringent DELTA-BLAST analyses for other pandoraviruses (e-value ≤ 1e-3 and identity ≥ 30% as thresholds) showed that 12 predicted translated ORFs had a hit against a domain of an enzyme involved in the TCA cycle, which was confirmed by BLASTp analysis against the nr database. These 12 ORFs putatively encode an acetyl-coenzyme A synthetase, a citrate synthase, an aconitase, a NADP-dependent IDH, a succinate dehydrogenase, and a malate dehydrogenase (Table 2). Moreover, DELTA-BLAST analysis revealed that a single translated ORF from *P. neocaledonia* (YP_009482013.1) harbored similarity to NADH dehydrogenase, an enzyme involved in the respiratory chain. This result was confirmed by BLASTp analysis against the nr database (bitscore: 45; identity: 31% with PKL55719.1, NADH:ubiquinone oxidoreductase from Methanomicrobiales archaeon HGW-Methanomicrobiales-6).

**Table 2:**
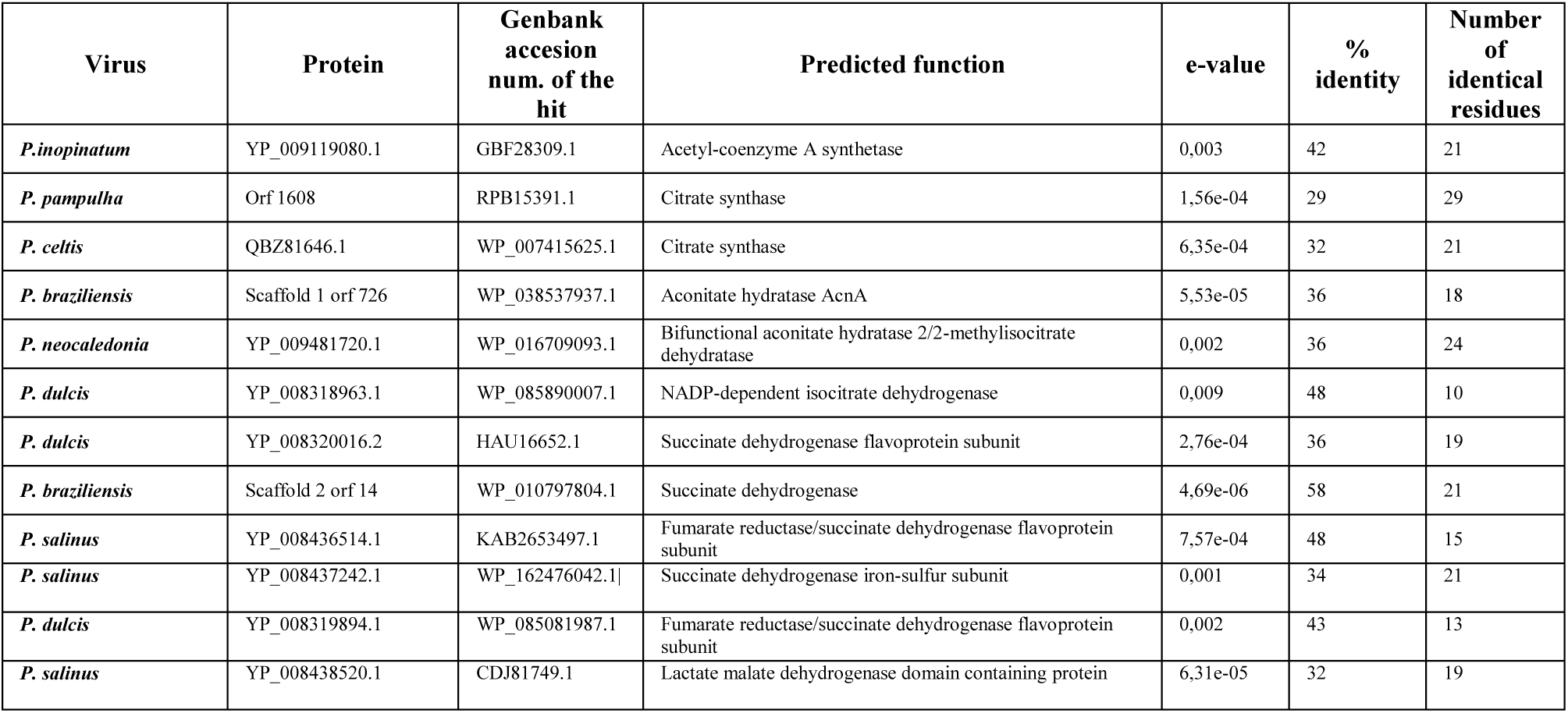
Results of the DELTA-Blast analyses carried out on the pandoraviruses.

BLASTp analysis against the COG database provided a hit for COG0277 (FAD/FMN-containing dehydrogenase) in all but two (*P. celtis* and *P. macleodensis*) pandoraviruses, with e-values ranging from 9.5e-68 and 9.1e-62 and identity percentages between 30.6 and 33.5% for alignment lengths ranging from 514 to 532 amino acids.

A hit was also found in *P. salinus, P. dulcis, P. inopinatum* and *P. pampulha* for COG1254 (acylphosphatase), with e-values ranging from 2.1e-15 to 2.57e-12 and identity percentages from 23.5 to 28.2% for alignments ranging from 156 to 239 amino acids in length.

### Transcriptomics of *P. massiliensis*: RNA-seq and qRT-PCR

RNA sequencing revealed that 6 of the 8 predicted *P. massiliensis* ORFs were transcribed at different time points of the viral cycle, especially between H4 and H8 post-infection (figure 8). No transcripts were detected for ORF595 (putative acetyl-coenzyme A synthetase) or ORF181 (putative succinate dehydrogenase). qRT-PCR revealed that 8 ORFs were transcribed at different time points of the viral cycle. For ORFs 595, 577, 1245, 132, 864, 762 and 206, the lowest Ct values were globally found between H8 and H16 post-infection. Of note, ORF181, the gene product of which exhibited a low similarity to succinate dehydrogenase, was transcribed at only one time point: 16 h post-infection (Table 3).

**Table 3:** Detection of *P. massiliensis* predicted TCA ORFs by qRT-PCR at different time points.

| ORFs | Predicted Enzyme | H0 | H2 | H4 | H6 | H8 | H10 | H12 | H16 |
| --- | --- | --- | --- | --- | --- | --- | --- | --- | --- |
| ORF595 | Acetyl CoA synthetase | NA | NA | NA | NA | 37 | NA | 31 | 30 |
| ORF577 | Citrate synthetase | NA | 35 | 34 | 36 | 38 | NA | 30 | 28 |
| ORF1245 | Aconitase | NA | 34 | 31 | 30 | 34 | NA | 25 | 27 |
| ORF132 | Isocitrate dehydrogenase | NA | NA | 38 | 38 | 38 | NA | 24 | 25 |
| ORF864 | Isocitrate dehydrogenase | NA | NA | 37 | 35 | 31 | NA | 32 | 30 |
| ORF762 | α-ketoglutarate decarboxylase | NA | NA | NA | 36 | 27 | NA | 22 | 23 |
| ORF181 | Succinate dehydrogenase | NA | NA | NA | NA | NA | NA | NA | 28 |
| ORF206 | Fumarase | NA | NA | 31 | 29 | 27 | 31 | 25 | 25 |

Footnotes: The numbers in each case are the Ct obtained for qPCR. NA: No amplification.

### Proteome Analysis of *P. massiliensis*

Proteomics analysis allowed us to identify 182 proteins, of which 162 (89%) were found in mature virions and 20 (11%) during the viral cycle. The function of most of these proteins is unknown. Two *P. massiliensis* proteins predicted to be involved in an energy production pathway were identified by this analysis in mature particles: ORF762 (putative α-ketoglutarate decarboxylase) and ORF595 (putative acetyl-coenzyme A synthetase) with an identity percentage of 100%.

### Functional tests of enzymatic function

Genes encoding the predicted enzymes of interest were synthesized and transferred to competent *E. coli* for recombinant protein expression. Soluble proteins were obtained for ORFs 577 (citrate synthase), 1245 (aconitate hydratase), 132, 864 (IDH), 787 and 1146 (α-ketoglutarate decarboxylases). We investigated the potential IDH activity of ORFs 132 and 864. We could determine a specific activity of 4 mU/mg for ORF 132, but no activity for ORF 864 (Figure 9). Kinetic assays were also performed to evaluate the catalytic parameters of ORF132. According to Michaelis-Menten equation fitting (R2=0.993), the following parameters were estimated: kcat=6.8×10-4 s-1, Km=51.8 µM and kcat/Km=13.12 s-1.M-1. In parallel, we performed the same analysis of human IDH from Sigma-Aldrich (St. Louis, MS, USA) and could determine a specific activity of 6.3 U/mg using the previously mentioned kit, as well as a kcat=16.3 s-1, Km=585.4 µM and kcat/Km=2.78×104 s-1.M-1 with an R2 of 0.997.

**Figure 9:**
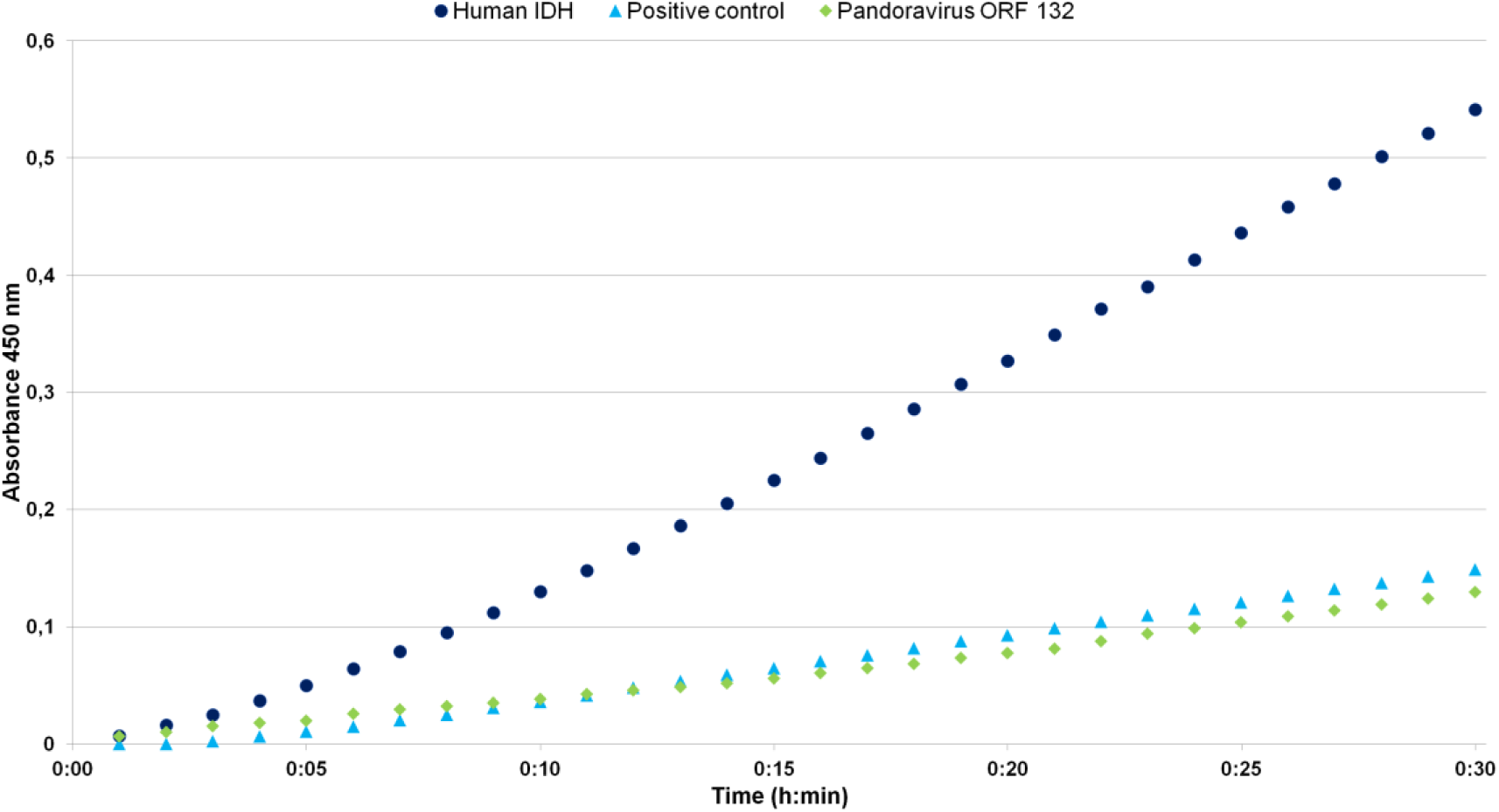
Evaluation of the enzymatic IDH activity of *P. massiliensis* ORF132.

Other predicted activities were also tested for ORFs 577, 1245, 787 and 1146 using the Citrate Synthase Assay kit, Aconitase Activity Assay kit and α-ketoglutarate Dehydrogenase Activity Colorimetric Assay kit from Sigma-Aldrich (St. Louis, MS, USA). However, no activity could be observed for these enzymes (data not shown).

## Discussion

In the present study, we identified virion membrane potential for the first time using two different fluorescent mitochondrial dyes, MitoTracker Deep Red 633 and TMRM, which allowed the detection of a fluorescent signal in mature virions of *P. massiliensis*. TMRM has been scientifically acknowledged as the best marker to assess mitochondrial membrane potential (28). For each experiment, critical negative controls were used to avoid false positive results. The intensity of this membrane potential was abolished following treatment with CCCP, a decoupling agent, confirming the accuracy of our observation. These intriguing findings represent the first experimental observation that a virus can have a membrane voltage. The search for predicted *P. massiliensis* proteins potentially involved in energy metabolism was unsuccessful for all enzymes except those in the TCA cycle. Of these enzymes, IDH was observed to be functional. The results showing that the virion membrane potential could be modified following addition of variable concentrations of acetyl-CoA, a known regulator of the TCA cycle, confirmed our findings. Indeed, bioinformatics analyses showed that *P. massiliensis* possessed all the genes encoding for enzymes of the TCA cycle (also called the Krebs or citric acid cycle). However, bioinformatics analysis revealed low sequence similarities with *bona fide* TCA orthologs. Furthermore, RNA-seq and confirmation by RT-PCR demonstrated that the predicted *P. massiliensis* TCA ORFs were all transcribed at the same time points, especially at the end of the developmental cycle of the virus (Figure 8). Two products of these genes were identified by proteomic analyses in mature particles, and the product of at least one predicted ORF, ORF132 encoding IDH, was shown to be functional. In the TCA cycle, IDH converts the isocitrate in α-ketoglutarate in the presence of the NAD+ or NADP+ cofactor. In nature, IDH catalyzes a catabolic reaction, during which NAD+ abstracts a hydride ion, which is a highly stereospecific enzymatic mechanism. This functionality cannot simply be the result of chance. The IDH step of the TCA cycle is often an irreversible reaction, with an overall estimated free energy of −8.4 kJ/mol (29). It is regulated by substrate availability, product inhibition, and competitive feedback inhibition by ATP (30). Isocitrate binds within the enzyme active site, which is composed of 8 amino acids. The metal ion Mg^2+^ or Mn^2+^ binds to three conserved Arg residues through hydrogen bond networks. The cofactor NAD^+^ or NADP^+^ binds within four regions with similar properties among the IDH enzymes, located around amino acids [250–260], [280–290], [300–330], and [365–380] (31). ORF132 is 146 amino acids long, while the known IDH from *E*.*coli* is 417 long (QJZ24410.1). IDH is typically multimeric (32) suggesting that the pandoraviral form may also form multimers. If the pandoraviral IDH is an ancestral form of the current eukaryotic IDH, then it is not surprising that the Km and Kcat of ORF132 are low. Subsequently, enzymes involved in the TCA cycle have evolved towards optimal performance with higher Km and Kcat values.

The TCA cycle is the central metabolic hub of cells. It is an exergonic catabolic energy acquisition pathway, which results in the oxidation of an acetyl group (derived from carbon compounds) to two molecules of carbon dioxide with the concomitant harvesting of high-energy electrons. Those electrons generate a proton gradient across the inner mitochondrial membrane through oxidative phosphorylation, with the aim of producing ATP through ATP synthase (33). The TCA cycle also provides, among other things, oxaloacetate for gluconeogenesis, intermediates for amino acid biosynthesis, nucleotide bases, cholesterol and porphyrins (33, 34). Of note, not all eukaryotes have mitochondria (35, 36) and the TCA cycle can occur in anaerobic organisms through the use of fumarate, nitrate, or various other compounds as terminal electron acceptors instead of O_2_ (37, 38).

As *P. massiliensis* is neither a eubacterium nor a eukaryote, the role of the TCA cycle and the existence of a membrane potential in mature particles is currently enigmatic. In eukaryotic mitochondria and bacteria, the membrane potential allows cells to function as a battery and generate energy. In eukaryotic cells, mitochondrial membrane potential results in the production of ATP via the TCA cycle. For *P. massiliensis*, we could not detect the production of ATP in mature particles (unpublished data). We could only observe a lower number of viral particles on amoeba cells infected with virions preincubated with CCCP at H0 and H3 post-infection than in negative controls, which suggested that the membrane voltage might be involved in the infection process of amoeba cell particularly in the early stages of infection. Until now, few viral genes in gene viruses have been described as possibly involved in metabolic pathways such as fermentation, sphingolipid biosynthesis and nitrogen metabolism (14). *Ostreococcus tauri* virus encodes an ammonium transporter, which enables host growth rescue when cultured with ammonium as the sole nitrogen source (39). TetV-1 encodes a mannitol metabolism enzyme, a saccharide degradation enzyme as well as other key fermentation genes (40). In all these previous cases, the viral genes seem to be host-derived and considered to be involved in viral manipulation of the host metabolism. It has also been shown that some giant viruses, including pandoraviruses, harbor cytochrome P450 genes, encoding enzymes that are known to be essential in the metabolism of endogenous regulatory molecules and exogenous drugs, but not their ancillary enzymatic redox partners, which could be recruited from host (12). Recently, a deep analysis of 501 environmental metagenome-assembled genomes of NCLDV revealed a diversity of metabolic genes involved in nutrient uptake, light harvesting, nitrogen metabolism, glycolysis and the TCA cycle (41). Moreover, tupanviruses possess a gene encoding citrate synthase, the first enzyme in the TCA cycle, for which no homologs were found in any other known virus. Phylogenetic analyses showed an independent origin of this gene in tupanviruses, which may have been acquired by tupanviruses via horizontal gene transfer from sympatric bacteria.

The phylogenetic origin of the TCA cycle may be the reverse TCA cycle, an endergonic anabolic pathway [37] that is used by some bacteria to produce carbon compounds [37-43]. However, the evolutionary history of the TCA cycle has not been completely elucidated, though it has been suggested that prior to endosymbiotic events, this pathway operated only as isolated steps [44]. Thus, the origin of its enzymes might be associated with lateral gene transfer or duplication events, suggesting other possible functionalities than those currently known [45]. Our data indicate that new candidates must be considered in the search for the origin of this cycle.

Evidence of energy production and the finding that a virus can carry enzymes encoding a TCA cycle opens a new horizon in giant virus research. Further experiments investigating the crystal structures of the viral enzymes, especially those of IDH, will be of particular interest for advances in the comprehension of this mysterious pandoravirus. Moreover, the use of yeast and bacterial complementation experiments to confirm the predicted enzymatic functions seems to be the logical next steps of this work. The database of newly discovered giant viruses is constantly growing and constitutes an important resource for the search for similar predicted enzymatic functions potentially involved in metabolic pathways.

## Conclusions

*P. massiliensis* undermines the last known historical viral hallmark, the lack of the Lipman system. Thus, the findings presented herein raise questions concerning whether pandoraviruses can still be classified biologically as a virus and renews arguments regarding the living nature of viruses in general.

